# Visualizing PINK1 Activity Dynamics in Single Cells with a Phase Separation-Based Kinase Activity Reporter

**DOI:** 10.1101/2025.10.08.681260

**Authors:** Katie G. Vineall, Alexia Andrikopoulos, Michael J. Sun, Anna Yan, Ethan R. Hartanto, Danielle L. Schmitt

**Affiliations:** Department of Chemistry and Biochemistry, University of California, Los Angeles, Los Angeles, CA USA; Institute for Quantitative and Computational Biosciences, University of California, Los Angeles, Los Angeles, CA USA; Molecular Biology Institute, University of California, Los Angeles, Los Angeles, CA USA

**Keywords:** PINK1, biosensor, fluorescence, mitophagy, kinase activity reporter

## Abstract

Phosphatase and tensin homologue-induced kinase 1 (PINK1) is a serine/threonine kinase that plays roles in mitophagy, cell death, and regulation of cellular bioenergetics. Current approaches for studying PINK1 function depend on bulk techniques that can only provide snapshots of activity and could miss the dynamics and cell-to-cell heterogeneity of PINK1 activity. Therefore, we sought to develop a novel PINK1 kinase activity reporter to characterize PINK1 activity. Taking advantage of the separation of phases-based activity reporter of kinase (SPARK) design, we developed a phase separation-based PINK1 biosensor (PINK1-SPARK). With PINK1-SPARK, we observe real-time PINK1 activity in single cells treated with mitochondria depolarizing agents or pharmacological activators. We then developed a Halo Tag-based PINK1-SPARK for multiplexed imaging of PINK1 activity with live-cell markers of mitochondrial damage. Thus, PINK1-SPARK is a new tool that enables temporal measurement of PINK1 activity in single live cells, allowing for further elucidation of the role of PINK1 in mitophagy and cell function.

## Introduction

Phosphatase and tensin homologue-induced kinase 1 (PINK1) is a mitochondrial serine/threonine kinase involved in mitophagy, the selective degradation of damaged mitochondria. In healthy mitochondria, PINK1 is constitutively recruited to the outer mitochondrial membrane and imported into the inner membrane space, where it undergoes N-terminal cleavage by proteases and subsequent degradation in the cytosol^1^. Following mitochondrial damage, the inner mitochondrial membrane becomes depolarized, and PINK1 is stabilized in its full-length form. This results in the dimerization, trans-autophosphorylation, and activation of PINK1, allowing it to phosphorylate and activate the E3 ubiquitin ligase Parkin and ubiquitin^2^. Parkin catalyzes the ubiquitination of outer mitochondrial membrane proteins, tagging damaged mitochondria for degradation by autophagic machinery^3,4^. Loss of function, autosomal recessive mutations in PINK1 are known to cause early onset Parkinson’s disease (EOPD), leading to the onset of symptoms, including tremor, rigidity, and bradykinesia, at a mean age of 31 years old^5,6^. While advances have been made in understanding the role of PINK1 in mitochondrial dynamics and the development of Parkinson’s disease, the current tools available to study PINK1 only provide snapshots of PINK1 activity and cannot distinguish PINK1 activity at the single cell level. We therefore sought to develop a kinase activity reporter capable of temporally characterizing PINK1 activity dynamics in live cells.

Fluorescent protein-based kinase activity reporters (KARs) offer a unique method to study kinase activity dynamics in single cells with high spatiotemporal resolution. KARs have been developed for several kinases including AMP-activated protein kinase (AMPK), protein kinase A (PKA), and protein kinase C (PKC)^7^. A recent KAR design, separation of phases-based activity reporter of kinase (SPARK), takes advantage of liquid-liquid phase separation for the detection of kinase activity^8^. In this KAR, phosphorylation of the SPARK construct increases the number of fluorophores per pixel, resulting in a simple readout of kinase activity that can be multiplexed with other biosensors^9^. SPARK-based biosensors have been made for kinases including PKA, ataxia-telangiectasia mutated (ATM), and AMPK^8–10^. As the SPARK design enables both time-lapse and snapshots of kinase activity in single cells, we sought to design a SPARK-based KAR for PINK1.

Here, we report the development of PINK1-SPARK utilizing the PINK1 phosphomotif from ubiquitin, a canonical PINK1 substrate. We find that in a variety of cell types expressing the biosensor, PINK1-SPARK robustly reports PINK1 activity following mitochondrial depolarization or direct activation of PINK1. We further engineer PINK1-SPARK to use the self-labeling protein HaloTag as a reporting unit for multiplexed imaging of PINK1 activity and reporters of mitochondrial depolarization. Altogether, we present a novel tool to measure PINK1 activity in single live cells, advancing the toolbox for studying this critical regulator of mitochondrial function.

## Results

### Development and characterization of PINK1-SPARK

PINK1-SPARK is comprised of two components - the first consists of a PINK1-specific substrate peptide sequence fused to enhanced green fluorescent protein (EGFP) and a hexameric coiled coil domain (HOTag3). The second component is made up of a phosphoamino acid binding domain fused to a tetrameric coiled coil domain (HOTag6). For approximately equal expression of both sensor components from the same plasmid, a bicistronic vector with a T2A ribosomal skip site sequence was used between the two components. In this design, activation of PINK1 would lead to multivalent protein-protein interactions and the visualization of green fluorescent droplets (**Fig 1A**).

**Figure 1.**
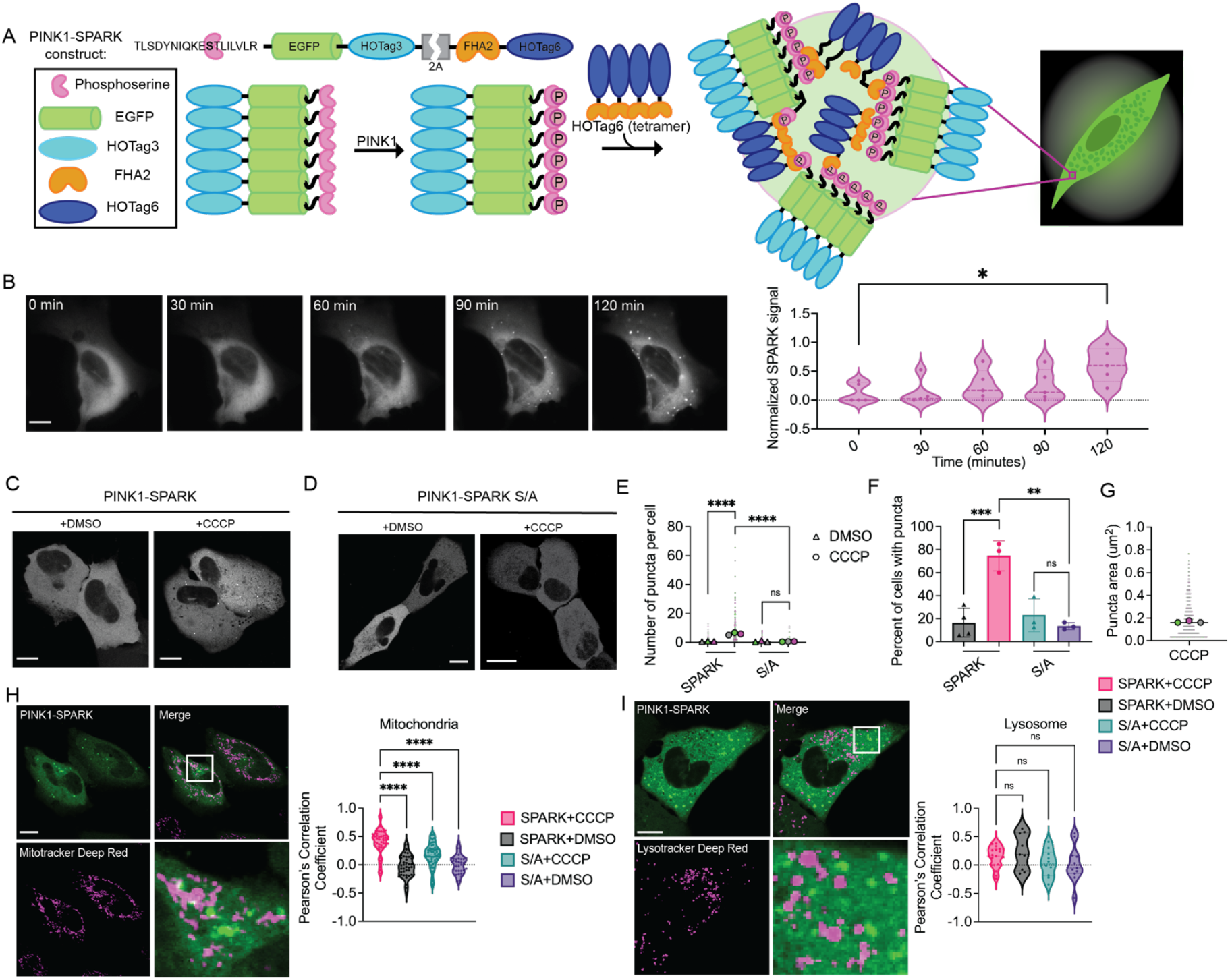
Design, development, and characterization of PINK1-SPARK. A. General scheme of reporter function. The first component consists of the PINK1-specific ubiquitin consensus sequence, EGFP, and the hexameric homooligomeric tag, HOTag3. The phosphoamino acid binding domain, FHA2, and the tetrameric HOTag6 make up the second component. PINK1 phosphorylates Ser65 within its consensus sequence, allowing recognition and binding by FHA2. The resulting proximity of multivalent interactions promotes phase separation and allows for the visualization of PINK1 activity as green fluorescent droplets. B. Left: time lapse images of U2OS cells transiently expressing PINK1-SPARK following addition of 10 µM CCCP. Middle: normalized PINK1-SPARK signal plotted against time. Right: violin plots of normalized SPARK signal at various time points (n = 5 cells from 2 experiments, *p = 0.0137, unpaired t-test). C. Representative images of U2OS cells expressing PINK1-SPARK following treatment with DMSO or 10 µM CCCP. D. Representative images of U2OS cells expressing the phosphonull PINK1-SPARK containing an S to A mutation (S/A). E. Number of puncta per cell in U2OS cells expressing PINK1-SPARK treated with DMSO (n = 193 cells from 3 experiments) or 10 µM CCCP (n = 226 cells from 3 experiments) or the phosphonull mutant following treatment with DMSO (n = 158 cells from 3 experiments) or 10 µM CCCP (n = 145 cells from 3 experiments) (****p < 0.0001, ordinary one-way ANOVA). F. Percent of cells showing puncta from figures C and D (black = PINK1-SPARK+DMSO; pink = PINK1-SPARK+CCCP; teal = S/A+CCCP; purple = S/A+DMSO; ***p = 0.0003, ordinary one-way ANOVA). G. Puncta area when observed following treatment with DMSO or 10 µM CCCP (ns = 0.0634, unpaired t-test, two-tailed). H. Left: representative image of U2OS cells expressing PINK1-SPARK stained with Mitotracker Deep Red. Right: colocalization analysis of PINK1-SPARK signal with mitochondria as quantified by Pearson’s correlation coefficient. U2OS cells expressing PINK1-SPARK treated with 10 µM CCCP (pink, n = 49 cells from 3 experiments) or DMSO (black, n = 39 cells from 3 experiments). Cells expressing the phosphonull mutant treated with 10 µM CCCP (blue, n = 55 cells from 3 experiments) or DMSO (purple, n = 33 cells from 3 experiments). (****p<0.0001, ordinary one-way ANOVA). I. Left: representative images of U2OS cell expressing PINK1-SPARK stained with Lysotracker Deep Red. Right: colocalization analysis of PINK1-SPARK signal with lysosomes. Cells expressing PINK1-SPARK treated with 10 µM CCCP (pink, n = 21 cells from 2 experiments) or DMSO (black, n = 9 cells from 2 experiments). Cells expressing the phosphonull mutant treated with 10 µM CCCP (blue, n = 10 cells from 2 experiments) or DMSO (purple, n = 11 cells from 2 experiments) (ns = 0.1861, ordinary one-way ANOVA). For all images, scale bars represent 10 µm. Bar graphs display mean ± standard deviation.

To achieve visualization of PINK1 activity at the single cell level, we first created four prototype PINK1-SPARKs with varying PINK1 phosphorylation motifs and phosphoamino acid binding domains (**Supplemental Figure 1A**). To assess the response of each candidate reporter, each PINK1-SPARK prototype was expressed in U2OS osteosarcoma cells and treated with carbonyl cyanide m-chlorophenylhydrazone (CCCP), an uncoupler of oxidative phosphorylation. The change in number of cells containing puncta was quantified before and after CCCP addition. All candidates showed an increase in the number of cells with puncta; however, PINK1-SPARK version 1.2 showed the greatest increase (68 ± 5.56%) following CCCP addition (**Supplemental Figure 1B**). PINK1-SPARK 1.2 contains the S68 phosphorylation motif from ubiquitin with an H-to-I mutation in the +3 position from the phosphosite to better recruit forkhead-associated domain 2 (FHA2) binding upon PINK1 phosphorylation^11^. Time-lapse imaging of PINK1-SPARK revealed the formation of puncta over 120 minutes following CCCP addition and an increased normalized SPARK signal over time (**Fig 1B**), which we corroborated with western blotting for phosphoubiquitin (**Supplemental Figure 1C**). We therefore moved forward with this variant, which we refer to as PINK1-SPARK, for further validation and characterization.

To ensure that the construct was not being phosphorylated at off-target sites, we developed a phosphonull PINK1-SPARK that contains a serine to alanine (S/A) mutation at the PINK1 phosphorylation site. Following exposure to CCCP, minimal puncta formation was measured in phosphonull PINK1-SPARK-expressing U2OS cells compared to cells expressing PINK1-SPARK (**Fig 1C-E**). Furthermore, a robust increase in the percent of cells with puncta was observed upon CCCP addition compared to S/A mutant PINK1-SPARK (74.6 ± 13.02%; **Fig 1F**).

We next sought to characterize PINK1-SPARK puncta. In U2OS cells, CCCP-induced PINK1-SPARK puncta had an average size of 0.165 ± 0.148 µm^2^ (**Fig 1G**). To determine if PINK1-SPARK puncta associate with organelles, U2OS cells expressing PINK1-SPARK were treated with CCCP and incubated with Mitotracker or Lysotracker Deep Red (**Fig 1H, I**). Quantitative colocalization analysis of SPARK signal and mitochondria revealed that PINK1-SPARK puncta had a significantly higher Pearson’s correlation coefficient than the DMSO-treated condition or cells expressing the phosphonull construct (DMSO: PCC = -0.0246 ± 0.1954; CCCP: PCC = 0.4004 ± 0.2253; ****p < 0.0001; **Fig 1H**). When colocalization analysis of PINK1-SPARK puncta and lysosomes was performed, no significant difference in Pearson’s correlation coefficient was observed between any condition (DMSO: PCC = 0.2356 ± 0.3136; CCCP: PCC = 0.1462 ± 0.1858; ns = 0.3371; **Fig 1I**), suggesting that PINK1-SPARK signal was not associated with lysosomes. As some Mitotracker dyes depend on intact mitochondrial membrane potential for accurate staining^12^, we also performed colocalization analysis between PINK1-SPARK and mitochondria labeled with mCherry-Cox8. When treated with CCCP, cells co-expressing PINK1-SPARK and mCherry-Cox8 showed a higher average Pearson’s correlation coefficient than the DMSO-treated condition, corroborating the results observed with Mitotracker (PCC = 0.510 ± 0.191); **Supplementary Figure 1D**).

As PINK1-SPARK relies on the formation of puncta driven by homooligomeric domains, we wanted to ensure PINK1-SPARK was not being recruited to stress granules. We immunostained U2OS cells expressing PINK1-SPARK treated with CCCP for fragile X messenger ribonucleoprotein (FMRP), a stress granule marker^13^ (**Supplemental figure 1E**). Colocalization analysis revealed no significant difference in PINK1-SPARK puncta association with stress granules when treated with CCCP (PCC = 0.497 ± 0.1306) or DMSO (PCC = 0.419 ± 0.146).

Finally, we estimated the Z-factor, a measure of assay fitness^14^. From the average response of PINK1-SPARK, we calculated the Z-factor to be 0.60, indicating good assay fitness. Thus, we have developed PINK1-SPARK, a robust reporter for PINK1 activity.

### PINK1-SPARK measures PINK1 activity in multiple cell types

We next used PINK1-SPARK to measure PINK1 activity in multiple cell lines, including HeLa, undifferentiated SH-SY5Y (USH), and neuronally differentiated SH-SY5Y (DSH). Puncta were observed in each cell line expressing PINK1-SPARK when treated with CCCP, but not when cells were treated with DMSO or expressed the phosphonull construct (**Fig 2A**). We confirmed that following 2-hour treatment with CCCP, an increase in the number of puncta and percent of cells with puncta was seen following CCCP addition in all cell types (HeLa: DMSO: 10.85 ± 12.09; CCCP: 69.00 ± 9.802; USH: DMSO: 0 ± 0; CCCP: 49.17 ± 32.63) DSH: DMSO: 3.333 ± 5.774; CCCP: 32.90 ± 17.30; **Fig 2B, C, E, F, H, I**). Comparison of puncta size between cell types treated with CCCP revealed that USH and DSH cells had larger puncta than HeLa (HeLa: 0.1518 ± 0.1537 µm^2^; USH: 0.2340 ± 0.2688 µm^2^; DSH: 0.2633 ± 0.1911 µm^2^; **Fig 2D, G, J**). Together, these data demonstrate the ability of PINK1-SPARK to characterize PINK1 activity in a variety of cell lines.

**Figure 2.**
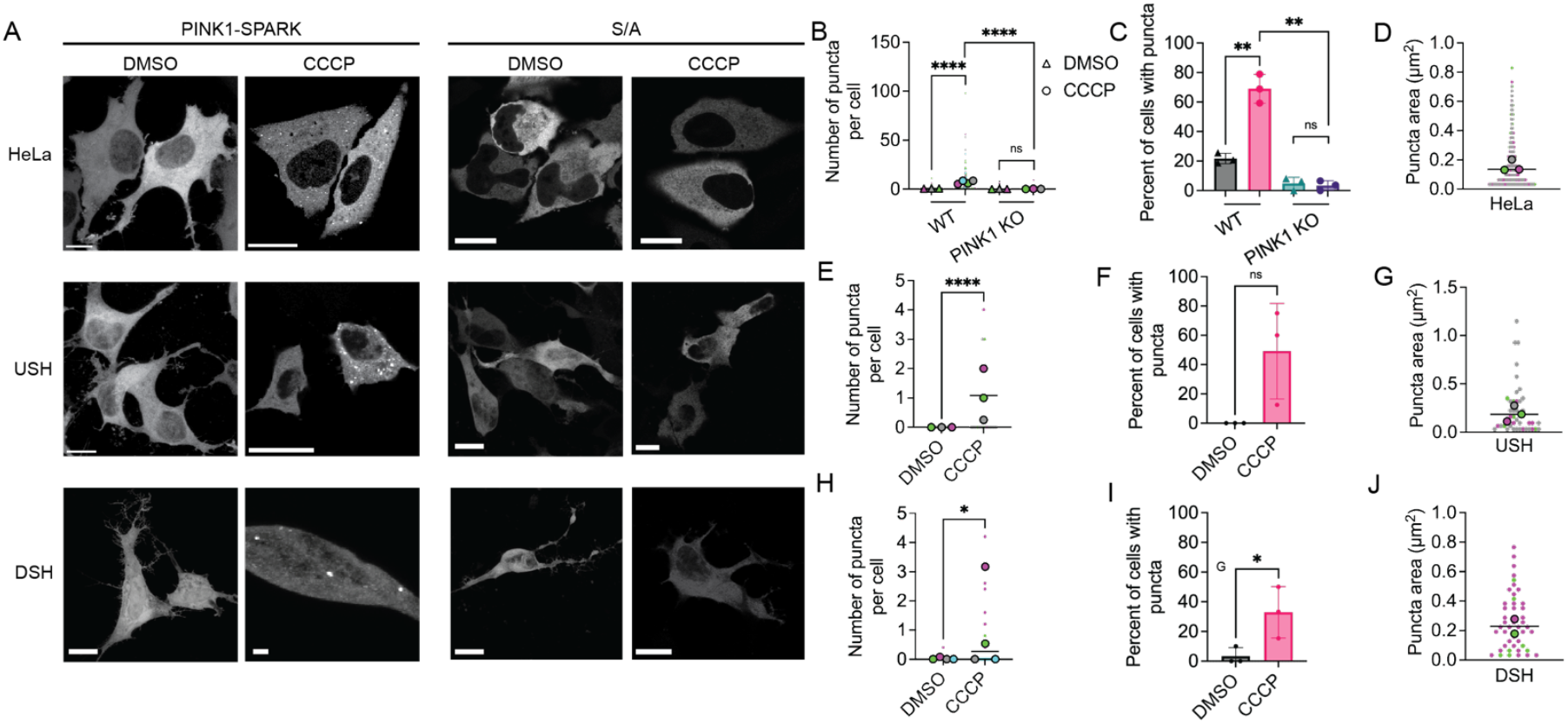
PINK1-SPARK characterizes PINK1 activity in multiple cell types. A. Left: representative images of HeLa, undifferentiated SH-SY5Y (USH), and neuronally differentiated SH-SY5Y (DSH) cells expressing PINK1-SPARK when treated with DMSO or 10 µM CCCP. Right: representative images of various cell types expressing the phosphonull mutant PINK1-SPARK (S/A) following addition of DMSO or CCCP. B. Number of puncta per cell in either WT HeLa or PINK1 KO HeLa cell following treatment with DMSO (WT: n = 141 cells from 3 experiments; PINK1 KO: n = 236 cells from 3 experiments) or 10 µM CCCP (WT: n = 159 cells from 4 experiments; PINK1 KO: n = 230 cells from 3 experiments; ****p < 0.0001; ns = 0.3962, unpaired t-test, two-tailed). C. Percentage of WT or PINK1 KO HeLa cells with puncta following addition of DMSO or 10 µM CCCP (**p = 0.0014; unpaired t-test, two-tailed). D. Puncta area in HeLa cells when treated with 10 µM CCCP (n = 463 puncta from 3 experiments). E. Number of puncta per USH cell following treatment with DMSO (n = 94 cells from 3 experiments) or 10 µM CCCP (n = 27 cells from 3 experiments; ****p < 0.0001; unpaired t-test, two-tailed). F. Percentage of USH cells with puncta following addition of DMSO or 10 µM CCCP (ns = 0.0594; unpaired t-test, two-tailed). G. Puncta area in USH cells when treated with 10 µM CCCP (n = 44 puncta from 3 experiments). H. Number of puncta per DSH cell following treatment with DMSO (n = 75 cells from 4 experiments) or 10 µM CCCP (n = 102 cells from 4 experiments; *p = 0.0278; unpaired t-test, two-tailed). I. Percentage of DSH cells with puncta following addition of DMSO or 10 µM CCCP (*p = 0.0484; unpaired t-test, two-tailed). J. Puncta area in DSH cells when treated with 10 µM CCCP (n = 48 puncta from 2 experiments). For all images, scale bars represent 10 µm. Bar graphs display mean ± standard deviation.

To confirm that PINK1-SPARK response is caused by the phosphorylation by PINK1 specifically, we used PINK1-SPARK in a PINK1 KO HeLa cell line^15^. PINK1 knock out was confirmed via Western blot (**Supplementary figure 2A**). Following treatment with CCCP, minimal puncta formation was seen (**Figure 2B, C**). Thus, PINK1-SPARK robustly and specifically reports PINK1 activity in live cells.

### PINK1-SPARK characterizes PINK1 activity in response to small molecule activators

PINK1 was previously considered undruggable, as until recently, there was minimal data on human PINK1 structure, and allosteric regulatory sites of PINK1 have not been identified. Despite these limitations, recent advances in the study of PINK1 biology have identified small molecule activators of PINK1, taking advantage of its potential for altered substrate specificity. Two such activators are N^6^-furfuryl ATP (kinetin triphosphate, KTP), an ATP substrate analog that is accepted by PINK1 with higher catalytic efficiency than ATP, and MTK458, a kinetin analog^16–18^.The metabolic precursor of KTP, kinetin riboside (KR), can be taken up by cells and converted to its nucleotide triphosphate form, which is reported to accelerate Parkin recruitment to depolarized mitochondria^17,18^. To measure PINK1 activity in single cells induced by KR, HeLa cells were treated with KR for 12 hours. We measured increased formation of PINK1-SPARK puncta compared to the DMSO-treated condition in WT HeLa (DMSO: 1.395 ± 4.559; KR: 88.05 ± 69.38), but not in PINK1 KO HeLa cells (DMSO: 0.1483 ± 0.8649; KR: 0.07101 ± 0.4574; **Fig 3A-B**). Next, we assessed PINK1 activation by MTK458 using PINK1-SPARK. While MTK458 is blood brain barrier-penetrable, it must be used in conjunction with a small concentration of CCCP that on its own does not induce significant mitophagy^16^. Following incubation with MTK458 and CCCP for 2 hours, we measured an increase in the number of puncta per cell compared to DMSO control and S/A mutant PINK1-SPARK in WT HeLa (DMSO: 0.6187 ± 1.481; MTK+CCCP: 22.34 ± 27.86; **Fig 3C-D**). No increase in puncta was observed following treatment with MTK458 and CCCP in PINK1 KO HeLa (DMSO: 0.1483 ± 0.8649; MTK+CCCP: 0.09950 ± 0.7550; **Fig 3D**). Finally, we compared puncta size between treatment conditions. WT HeLa cells treated with KR or MTK458 and CCCP had significantly larger puncta than cells treated with CCCP (KR: 0.1864 ± 0.1971 µm^2^; MTK+CCCP: 0.1919 ± 0.1681 µm^2^; CCCP: 0.1518 ± 0.1537 µm^2^; ***p = 0.0002; **Fig 3E**). Thus, PINK1-SPARK reports PINK1 activity induced by small molecules and corroborates the previous identification of KTP and MTK458 as activators of PINK1.

**Figure 3.**
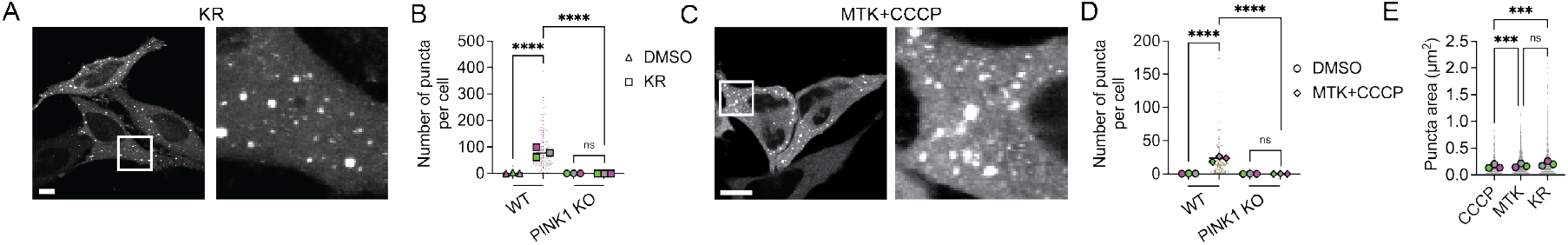
PINK1-SPARK visualizes PINK1 activity following multiple types of PINK1 activation. A. Representative image of HeLa cells expressing PINK1-SPARK when treated with 50 µM kinetin riboside (KR). B. Number of puncta per cell in WT or PINK1 KO HeLa expressing PINK1-SPARK treated with DMSO (WT: n = 118 cells from 3 experiments; PINK1 KO: n = 236 cells from 3 experiments) or 50 µM KR (WT: n = 129 cells from 3 experiments; PINK1 KO: n = 169 cells from 3 experiments; ****p < 0.0001; ns = 0.2897; unpaired t-test, two-tailed). C. Representative image of HeLa cells expressing PINK1-SPARK when treated with MTK458 (3.1 µM) and CCCP (0.5 µM). D. Number of puncta per cell in WT or PINK1 KO HeLa expressing PINK1-SPARK when treated with DMSO (circles, WT: n = 69 cells from 3 experiments; PINK1 KO: n= 236 cells from 3 experiments) or 3.1 µM MTK + 10 µM CCCP (diamonds, WT: n = 81 cells from 2 experiments; PINK1 KO: n = 201 cells from 3 experiments; ****p < 0.0001; unpaired t-test, two-tailed). E. Comparison of puncta size between treatment conditions: 10 µM CCCP (n = 481 puncta from 3 experiments), 3.1 µM MTK + 0.5 µM CCCP (n = 1451 puncta from 3 experiments), or 50 µM KR (n = 4727 puncta from 3 experiments; ***p = 0.0002; unpaired t-test, two-tailed). For all images, scale bars represent 10 µm. Bar graphs display mean ± standard deviation.

### Design of a PINK1-SPARK construct integrating HaloTag technology

To increase the multiplexing capabilities of SPARK technology, we developed a novel SPARK construct reliant upon the self-labeling protein HaloTag, termed Halo-PINK1-SPARK. Following activation of PINK1, cells expressing Halo-PINK1-SPARK are incubated with a deep red Janelia Fluor ligand (JF646), that covalently binds to HaloTag to fluorescently label PINK1-SPARK^19^ (**Fig 4A**). To validate the use of Halo-PINK1-SPARK, HeLa cells expressing Halo-PINK1-SPARK were incubated with DMSO or CCCP for 2 hours, then incubated with JF646. Similar to the original PINK1-SPARK construct, cells treated with CCCP showed an increased number of PINK1-SPARK puncta per cell and an increase in the percent of cells showing puncta compared to DMSO (CCCP: 69.00 ± 9.802%; DMSO: 21.70 ± 3.509%;) and phosphonull PINK1-SPARK (CCCP: 15.27 ± 7.838%; DMSO: 20.57 ± 5.095%; **Fig 4B-D)**. Interestingly, Halo-PINK1-SPARK showed a significantly increased puncta size compared to PINK1-SPARK (Halo-PINK1-SPARK: 0.2517 ± 0.3450 µm^2^; PINK1-SPARK: 0.1518 ± 0.1537 µm^2^; ****p < 0.0001; **Fig 4E**). To ensure that Halo-PINK1-SPARK was not being phosphorylated at off-target sites, we created a phosphonull (S/A) mutant construct that minimally formed puncta upon activation of PINK1 by CCCP (**Fig 4B-C**). Furthermore, phosphonull Halo-PINK1-SPARK did not show an increase in the percent of cells with puncta between DMSO or CCCP treatment (**Fig 4D**).

**Figure 4.**
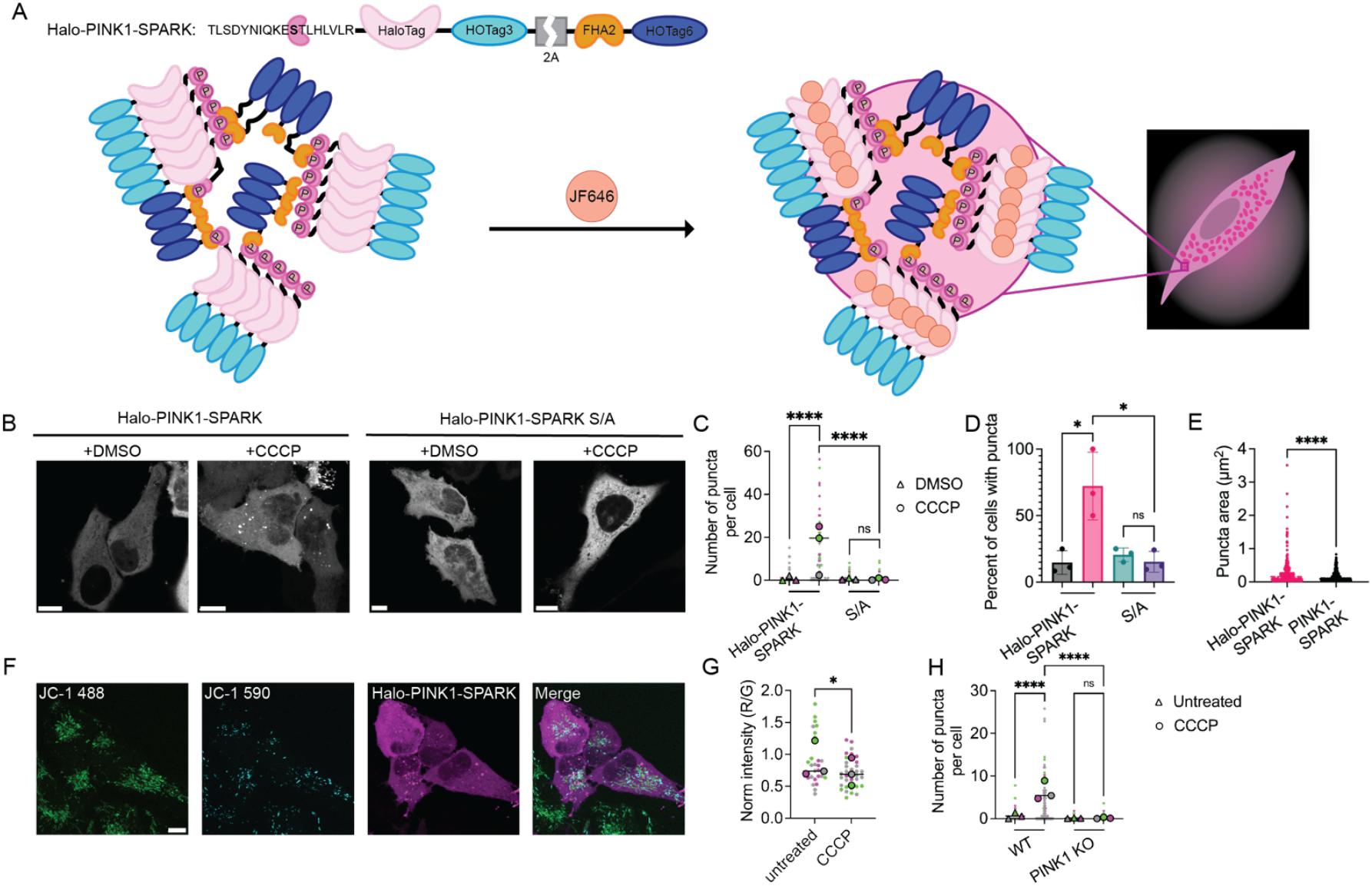
Halo-PINK1-SPARK allows for enhanced visualization of PINK1 activity dynamics and multiplexing with JC-1. A. General schematic of reporter function. Following activation of PINK1, cells are incubated for 30 minutes with 200 nM Janelia Fluor 646, fluorescently labeling Halo-PINK1-SPARK puncta. B. Representative images of HeLa cells expressing Halo-PINK1-SPARK or phosphonull Halo-PINK1-SPARK when treated with DMSO or 10 µM CCCP. C. Number of puncta per cell in HeLa expressing Halo-PINK1-SPARK or phosphonull Halo-PINK1-SPARK when treated with DMSO (Halo-PINK1-SPARK: n = 64 cells from 3 experiments; S/A: n = 73 cells from 3 experiments) or 10 µM CCCP (Halo-PINK1-SPARK: n = 59 cells from 3 experiments; S/A: n = 89 cells from 3 experiments; ****p < 0.0001; ns = 0.7156; unpaired t-test, two-tailed). D. Comparison of the percentage of HeLa cells with puncta following treatment with DMSO (black) or 10 µM CCCP (pink) (*p = 0.0477; ns = 0.3822; unpaired t-test, two-tailed). E. Comparison of Halo-PINK1-SPARK and PINK1-SPARK puncta size following 10 µM CCCP addition (****p < 0.0001; unpaired t-test, two-tailed). F. Representative images of HeLa cells transiently expressing Halo-PINK1-SPARK stained with JC-1 following treatment with 10 µM CCCP. G. Normalized fluorescence of JC-1 before and after treatment with 10 µM CCCP (untreated: n = 29 cells from 3 experiments; CCCP: n = 35 cells from 3 experiments; *p = 0.0239; unpaired t-test, two-tailed). H. Number of puncta per cell in WT or PINK1 KO HeLa before (triangles, WT: n = 88 cells from 3 experiments; PINK1 KO: n = 130 cells from 3 experiments) and after treatment with 10 µM CCCP (circles, WT: n = 111 cells from 3 experiments; PINK1 KO: n = 152 cells from 3 experiments; ****p < 0.0001; ns = 0.4948; unpaired t-test; two-tailed). For all images, scale bars represent 10 µm. Bar graphs display mean ± standard deviation.S

### Halo-PINK1-SPARK enables multiplexed imaging of PINK1 activity with markers of mitochondrial damage

To quantify PINK1 activity and mitochondrial depolarization, we performed end-point imaging of HeLa cells expressing Halo-PINK1-SPARK and treated with the mitophagy reporter dye JC-1. JC-1 is a cationic dye that aggregates in mitochondria, fluorescing red when the membrane potential is intact and fluorescing green when membranes become depolarized^20^. As Halo-PINK1-SPARK utilizes a deep red fluorophore, multiplexing of mitochondrial membrane potential and PINK1 activity can be performed. When treated with CCCP, we predicted that HeLa cells co-expressing Halo-PINK1-SPARK and JC-1 would show a decrease in normalized JC-1 fluorescence (RFP/GFP) alongside an increase in the number of puncta per cell due to PINK1 recruitment following depolarization of the mitochondrial membrane. When HeLa cells expressing PINK1-SPARK stained with JC-1 were incubated with 10 µM CCCP for 2 hours, a decrease in JC-1 signal and an increase in the number of puncta per cell were observed (untreated: 1.000 ± 2.642; CCCP: 6.375 ± 6.853 puncta per cell; ****p < 0.0001; **Fig 4F-H**). As a control, PINK1 KO HeLa cells transiently expressing Halo-PINK1-SPARK were stained with JC-1 and were observed to show significantly less puncta than WT HeLa in both untreated (0.1308 ± 0.5339) and CCCP-treated cells (0.1645 ± 0.8093; **Fig 4H**). Using HaloTag technology, we have enabled enhanced multiplexing capabilities using the SPARK construct with markers of mitochondrial damage.

## Discussion

Genetically encoded fluorescent protein-based kinase activity reporters have proven to be a useful tool in understanding the dynamics of signaling networks regulating metabolism and other cell functions. In the present study, we designed a kinase activity reporter based on the SPARK design for PINK1, termed PINK1-SPARK. While tools have been previously developed to study mitophagy flux, none are capable of directly reporting PINK1 activity^21–24^. For the first time, we are able to study PINK1 activity dynamics at the single-cell level using PINK1-SPARK, expanding the toolkit to study PINK1-mediated mitophagy and ultimately EOPD pathology. PINK1-SPARK has good assay fitness as measured by the Z-factor, making it a suitable assay for a wide variety of applications. While PINK1-SPARK has a large dynamic range, its spatial resolution falls short of that of FRET-based kinase activity reporters, which can measure kinase activity at distinct subcellular locations^7^. Additionally, we did not test the reversibility of PINK1-SPARK, as PINK1 inhibitors are non-specific^25^, thus PINK1-SPARK functions best as a “turn on” reporter of PINK1 activity. Overall, PINK1-SPARK represents a significant advancement in the detection and quantification of PINK1 activity.

Using PINK1-SPARK, we performed time lapse fluorescence imaging following addition of the mitochondrial uncoupler CCCP. We found that PINK1-SPARK showed increased puncta following CCCP addition, but not DMSO. Furthermore, the phosphonull PINK1-SPARK mutant showed no puncta formation, even in conditions of active PINK1, demonstrating that there is minimal off-target phosphorylation of the reporter. The lack of PINK1-SPARK response to CCCP incubation in PINK1 KO HeLa cells demonstrates the specificity of the reporter for PINK1. Colocalization analysis revealed that PINK1-SPARK signal is associated with mitochondria but not associated with lysosomes or stress granules. We further validated the use of PINK1-SPARK in multiple cell types and observed an increase in the number of puncta per cell following treatment with CCCP. Interestingly, it was observed that HeLa cells showed significantly more puncta per cell and a larger increase in the percentage of cells with puncta following CCCP addition than USH or DSH cells. This could be explained by the difference in activation mechanisms of PINK1 between cell types. While CCCP addition is thought to increase PINK1 activity in HeLa cells by inhibiting proteolytic cleavage of PINK1^1^, SHSY5Y cells are suggested to respond by increasing transcription of the gene encoding PINK1, requiring a longer incubation period with CCCP for maximal PINK1 activity to be measured^26^. Through these studies, we have highlighted PINK1-SPARK as a versatile tool to probe PINK1 activity dynamics.

Building on these findings, we investigated changes in PINK1 activity induced by previously identified small molecule activators of PINK1. We found that number of puncta per cell increased following treatment with direct PINK1 activators KR and MTK458. KR treatment caused the largest increase in percentage of cells with PINK1-SPARK puncta compared to other PINK1 activators, corroborating previous findings that KR is a potent PINK1 activator^17,18^. Furthermore, these data show a graded pattern between the number of puncta per cell and puncta size, suggesting that the different PINK1 activators have varying effects on PINK1 activity. While the discovery of small molecule regulators for PINK1 has been limited by the lack of tools available to study PINK1 activity, the simple readout of PINK1-SPARK makes this biosensor a good candidate for use in high-throughput drug screenings.

In order to enhance the multiplexing capabilities of the SPARK construct, we developed a PINK1-SPARK containing HaloTag (Halo-PINK1-SPARK), a small protein capable of covalently binding fluorescent ligands with varying excitation and emission spectra. We found that incubation of cells expressing Halo-PINK1-SPARK with JF646 allowed for visualization of puncta with deep red emission. Halo-PINK1-SPARK had similar response to CCCP as EGFP-PINK1-SPARK in WT and PINK1 KO cells, validating its use as a robust and specific reporter of PINK1 activity. Using Halo-PINK1-SPARK, we then probed the interplay of mitochondrial depolarization and PINK1 activity using the mitochondrial stain JC-1. We were able to co-image both PINK1 activity and a readout of mitochondrial membrane potential within the same cell. Thus, Halo-PINK1-SPARK can be used for multiplexed imaging with other biosensors or probes. While not tested here, Halo-PINK1-SPARK could be used to readout PINK1 activity using other HaloTag fluorophores to maximize multiplexing capabilities, enhancing the utility of PINK1-SPARK.

In summary, we have generated a phase separation-based PINK1 activity reporter that we have used to study PINK1 activity in multiple cell types, with different activators, and for multiplexed imaging. Using our reporter, the study of PINK1 biology in single cells is now possible, which will enhance our current understanding of mitochondrial homeostasis and PINK1 function.

## Materials and Methods

### Plasmids

Primers were obtained from IDT. PINK1-SPARK was adapted from PKA-SPARK. Initial designs were cloned using Gibson Assembly (Gibson Assembly HiFi Master Mix, Fisher Scientific Cat #A46628) to insert the ubiquitin consensus sequence and phosphoamino acid binding domains. AKAR1 (encoding 14-3-3t) was provided by Jin Zhang^27^ and the FHA2 binding domain was cloned from CKAR. PCR was performed with PKA-SPARK as template, with subsequent DpnI digestion (Thermo Scientific, Cat# FD1703) and PCR cleanup (Thermo Scientific, Cat# K310001). Resulting fragments were subject to Gibson Assembly. QuikChange was used to generate the phosphonull PINK1-SPARK mutant^28^. Successful clone generation was confirmed using whole plasmid sequencing (Plasmidsaurus).

To generate Halo-PINK1-SPARK, two gene fragments encoding the PINK1-SPARK construct with HaloTag substituted for EGFP were designed using IDT gBlock Gene Fragments. PCR was performed on the fragments and PINK1-SPARK as template, and Gibson Assembly was used to clone the construct. Successful clone generation was confirmed using whole plasmid sequencing (Plasmidsaurus).

mCherry-Mito-7 was a gift from Michael Davidson (Addgene plasmid # 55102 ; http://n2t.net/addgene:55102 ; RRID:Addgene_55102). pcDNA3-PKA-SPARK was a gift from Xiaokun Shu (Addgene plasmid # 106920 ; http://n2t.net/addgene:106920 ; RRID:Addgene_106920). CKAR was a gift from Alexandra Newton (Addgene plasmid # 14860 ; http://n2t.net/addgene:14860 ; RRID:Addgene_14860).

### Cell culture and transfection

U2OS, HeLa, and SHSY5Y cell lines were obtained from ATCC (Cat #HTB-96; CRM-CCL-2; CRL-2266). PINK1 KO HeLa cells were provided by Ming Guo^15^. U2OS, WT HeLa, and PINK1 KO HeLa were maintained at 37°C and 5% CO_2_ in Dulbecco’s modified Eagle medium (Thermo Scientific, Cat #10569010) and 100 U/mL penicillin-streptomycin (Pen/Strep, Fisher Scientific, Cat #15-140-122). SHSY5Y cells were maintained at 37°C and 5% CO_2_ in Eagle’s minimum essential medium (Sigma-Aldrich, Cat #M4526) and 100 U/mL penicillin-streptomycin. All cells were routinely checked for mycoplasma using NucBlue (Fisher Scientific Cat #R37605) staining and a PCR mycoplasma detection kit (Fisher Scientific, Cat #AAJ66117AMJ).

U2OS and HeLa cells were seeded at 3 × 10^5^ one day before transfection in 35mm glass bottom dishes (Cellvis, Cat #d35-14-1.5-n). The following day, transfection was carried out in reduced serum medium (Opti-MEM, Thermo Scientific, Cat #31985070) using Fugene 4K (U2OS) or Fugene HD (HeLa). SHSY5Y cells were seeded at 6 × 10^5^ one day before transfection and transfection was performed using Mirus TransIT-X2 (Fisher Scientific, Cat #MIR 6000).

### SHSY5Y differentiation

SHSY5Y cells were terminally differentiated following previously published protocols^29^. SHSY5Y cells were grown to confluence in 35mm glass bottom 6-well plates (Fisher Scientific, Cat #NC0452316) in EMEM containing 10 µM retinoic acid (RA, Sigma-Aldrich, Cat #R2625) for 3 days followed by removal of the RA-containing media and replaced with EMEM containing 80 nM 12-o-tetradecanoyl-phorbol-13-acetate (TPA, Millipore Sigma, Cat #5.00582.0001) for another 3 days of differentiation. Transfection was then performed as previously described using Mirus TransIT-X2.

### Time lapse fluorescence imaging and image analysis

Fluorescence microscopy time-lapse experiments were performed on a Nikon ECLIPSE Ti2 epifluorescence microscope equipped with a CHI60 Plan Fluor 40X Oil Immersion Objective Lens (N.A. 1.3, W.D. 0.2 mm, F.O.V. 25mm; Nikon), a Spectra III UV, V, B, C, T, Y, R, nIR light engine featuring 380/20, 475/28, and 575/25 LEDs, Custom Spectra III filter sets (440/510/575 and 390/475/555/635/747) mounted in Ti cube polychroic, a Kinetix 22 back-illuminated sCMOS camera (Photometrics), and a stage-top incubator set to 37°C (Tokai Hit). NIS-Elements software (Nikon) was used. Exposure times between 10-100ms were used. Imaging began before adding DMSO or CCCP (10 µM, Fisher Scientific, Cat #04-525-00) and continued for 2 hours.

PINK1 activity was quantified in time-lapse imaging as the number of PINK1-SPARK puncta over time and SPARK signal. SPARK signal was calculated using previously published protocols - the sum of the punctate pixel fluorescence intensity and the cell’s pixel intensity was quantified using the Analyze Particle function in Fiji. The ratio of puncta to cell intensity was then determined. The normalized SPARK value was calculated using SPARK values at different time points normalized to the peak SPARK value^8–10^. TrackMate (Fiji) was used to segment and track puncta over time^30,31^.

### Confocal imaging and image analysis

Confocal microscopy was used to take end-point images of cells expressing PINK1-SPARK on a Leica Stellaris 5 DMi8 Confocal Microscope equipped with a DMOD WLL (440 nm – 790 nm), HC PL APO 63x/1.40 oil immersion CS2 lens, and a Power HyD S detector. LasX software (Leica) was used to control the microscope. Laser intensities between 0.5-8% were used to image cells. Cells were treated with DMSO, CCCP, or MTK458 (3.1 µM, MedChem Express, Cat #HY-152943) and CCCP (0.5 µM) for 2 hours before imaging. For end-point imaging following KR treatment, cells were treated with KR (50 µM, MedChem Express, Cat # HY-101055) for 12 hours.

Max Z projections were generated from Z-stack confocal images. Trainable Weka Segmentation (Fiji) was used to segment puncta^32^. Puncta number and size were quantified using the Analyze Puncta function in Fiji.

For colocalization analysis with mitochondria and lysosomes, cells expressing PINK1-SPARK were stained with 25 nM MitoTracker Deep Red (Thermo Scientific Cat #M22426) or LysoTracker Deep Red (Thermo Scientific Cat #L12492) for 10 minutes. Max Z projections were acquired as previously described and colocalization was quantified using BIOP-JacoP (Fiji)^30^. For co-imaging of PINK1-SPARK and mCherry-Cox8, U2OS cells were co-transfected with PINK1-SPARK and mCherry-Cox8 rather than stained with MitoTracker Deep Red. For multiplexing experiments, cells expressing Halo-PINK1-SPARK were stained with 200 nM JC-1 (Thermo Fisher, Cat #T3168) for 15 minutes. The mean gray value was calculated for each channel using the analyze function in ImageJ and normalized intensities were calculated by dividing the mean gray value for the red channel by that of the green channel.

### Western blot analysis

Cells were lysed in mammalian protein extraction reagent containing protease and phosphatase inhibitors (M-PER; Thermo Scientific Cat #78501; #A32963; #A32957) and resolved in 4-15% Mini-PROTEAN gels (Bio-Rad, Cat #4568083). After transferring, PINK1 rabbit mAb (Cell Signaling Technology, Cat #69465), ubiquitin (P4D1) mouse mAb (Abcam, Cat #AB303664), and phospho-ubiquitin (Ser65; E2J6T) rabbit mAb (Cell Signaling Technology, Cat #62802) were used at 1:1000. HRP-Goat-anti-Mouse (Bio-Rad, Cat #1706516) and HRP-Goat-anti-Rabbit (Bio-Rad, Cat #1706515) antibodies were used at 1:10000. Cofilin (D3F9) rabbit mAb (Cell Signaling Technology, Cat #5175S) was used at 1:1000 as a loading control. Clarity Western ECL Substrate (Bio-Rad, Cat #1705060) was used for visualization. To analyze Western blot signal, blots were imaged using an Azure 600 (Azure Biosystems, Cat #AZI600-01).

### Immunostaining

HeLa cells were plated on coverslips (Fisher Scientific, Cat #12-550-123) in a 6-well plate and transfected with PINK1-SPARK. One day after transfection, cells were treated with CCCP or DMSO for 2 hours and then fixed in 4% (v/v) paraformaldehyde for 20 minutes at room temperature. Following fixation, cells were permeabilized in 1X Dulbecco’s phosphate-buffered saline (DPBS) with 0.1% Triton X-100 (cat X) and 5% BSA per well. FMRP (D14F4) rabbit mAb (Cell Signaling Technology, Cat #7104) was added to coverslips at 1:200 overnight at 4°C. The next day, cells were incubated with HRP-Goat-anti-Rabbit Alexa Fluor 633 (1:200; Thermo Scientific, Cat #A-21070) protected from light for 1 hour at room temperature prior to mounting with Prolong Glass Antifade Mountant with NucBlue Stain (Thermo Scientific, Cat #P36985).

### Statistics and reproducibility

Figure preparation and statistical analysis were performed using GraphPad Prism 10. For comparison of two parametric data sets, two-tailed unpaired t-test was used. When appropriate, ordinary one-way ANOVA or two-way ANOVA followed by multiple comparisons was done. Statistical significance was defined as p < 0.05 with a 95% confidence interval. The number of trials, number of independent experiments, technical replicates, and statistical tests used are reported in all figure legends. Where appropriate, data was plotted using the SuperPlots method^33^. All dot plots shown depict the mean ± standard deviation.

## Data Availability Statement

All data are available within the main manuscript and the Supporting Information.

## Author Information

### Authors

Katie G. Vineall—Department of Chemistry and Biochemistry, University of California Los Angeles, Los Angeles, California, 90095, United States of America

Alexia Andrikopoulos—Department of Chemistry and Biochemistry, University of California Los Angeles, Los Angeles, California, 90095, United States of America

Michael J. Sun—Department of Chemistry and Biochemistry, University of California Los Angeles, Los Angeles, California, 90095, United States of America

Anna Yan—Department of Chemistry and Biochemistry, University of California Los Angeles, Los Angeles, California, 90095, United States of America

Ethan R. Hartanto—Department of Chemistry and Biochemistry, University of California Los Angeles, Los Angeles, California, 90095, United States of America

## Author Contributions

D.L.S., K.G.V, and A.A. conceived and designed PINK1-SPARK. K.G.V. developed and tested PINK1-SPARK and performed all time course imaging. K.G.V., A.A., M.J.S, A.Y., and E.H. performed confocal imaging experiments. A.A. performed western blot experiments. K.G.V., M.J.S., and A.A. analyzed microscopy data. D.L.S. oversaw all experiments. K.G.V. and D.L.S. wrote the manuscript. K.G.V. and D.L.S. prepared the figures. All authors contributed to the final version of the manuscript.

## Notes

The authors declare they have no competing interests.

## Acknowledgements

We thank Jin Zhang and Ming Guo for sharing plasmids and cell lines, Jack Scully for his assistance with molecular cloning, and all members of the Schmitt Lab for their helpful discussion. This work was supported by the National Institutes of Health (1DP2GM154012 to D.L.S. and T32GM007185 to A.A. and K.G.V.).

## Notes

### Competing Interest Statement

The authors have declared no competing interest.

### Summary of Updates

This version of the manuscript has been revised to reflect new data acquired since the original posting.

