## Supplemental Figures for "Visualizing PINK1 Activity Dynamics in Single Cells with a Phase Separation-Based Kinase Activity Reporter"

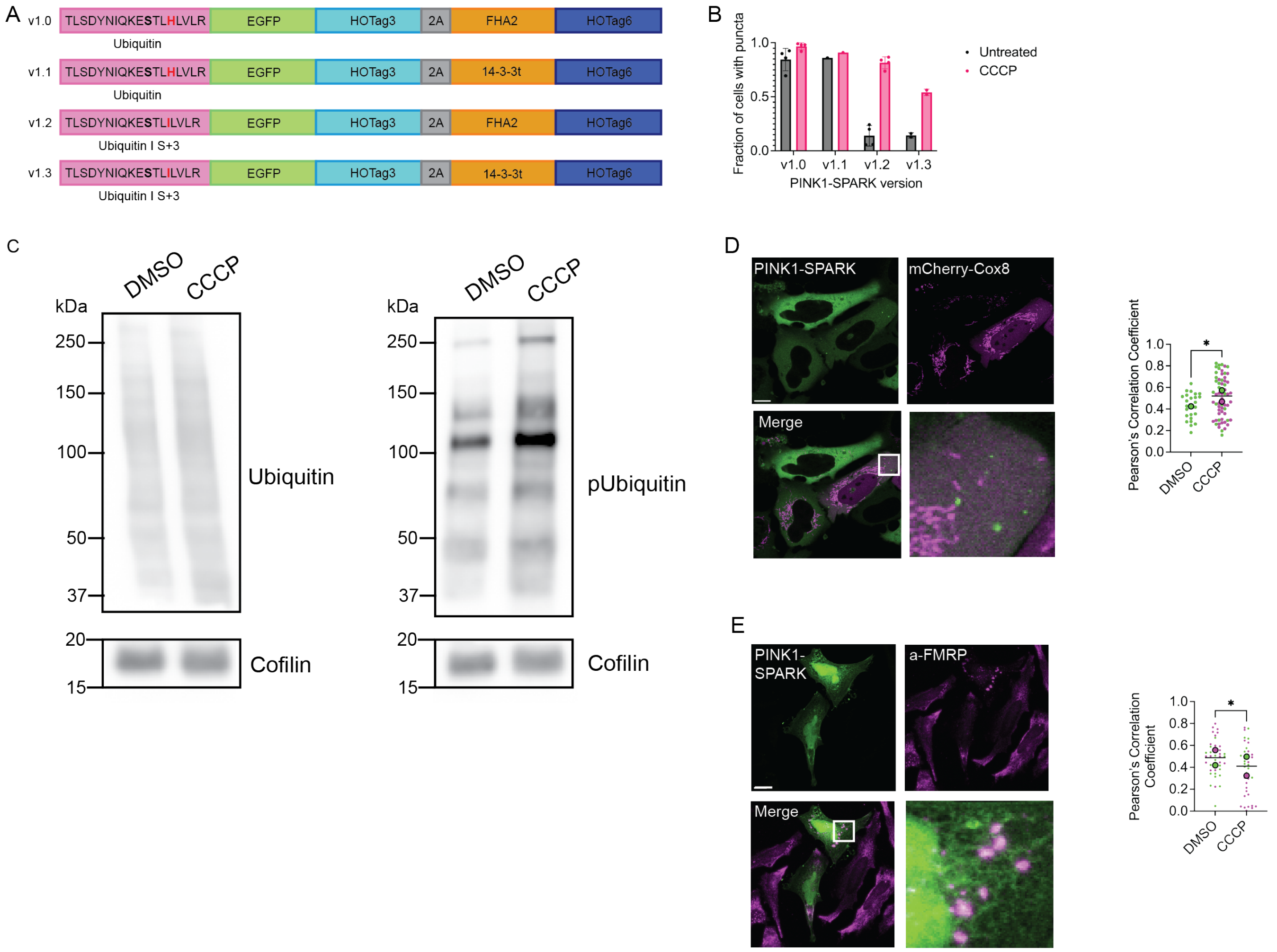


**Supplementary figure 1. Development and characterization of PINK1-SPARK v1.2.**

A. Design of the original PINK1-SPARK construct. PINK1-SPARK v1.0 is composed of the ubiquitin consensus sequence (S65 in bold) and the FHA2 phosphoamino acid binding domain. v1.1 consists of the ubiquitin consensus sequence and the 14-3-3t phosphoamino acid binding domain. v1.2 contains the ubiquitin consensus sequence with a mutation from H to I at the pS+3 position (red) and FHA2. v1.3 contains the mutated ubiquitin consensus sequence and 14-3-3t.
B. Percent of cells showing puncta before and after 10 µM CCCP treatment for each candidate reporter.
C. Western blot of ubiquitin and phosphoubiquitin levels in WT HeLa cells over 2 hours of treatment with DMSO or 10 µM CCCP, along with a cofilin loading control.
D. Left: representative images of U2OS cells co-expressing PINK1-SPARK and mCherry-Cox8. Right: colocalization analysis of PINK1-SPARK puncta and mCherry-Cox8 following incubation with DMSO or 10 µM CCCP (*p = 0.0419; unpaired t-test; two-tailed).
E. Left: immunofluorescence confocal images of HRP-Goat-anti-rabbit Alexa Fluor 488-labeled a-FMRP and Halo-PINK1-SPARK. Right: colocalization analysis of stress granule and Halo-PINK1-SPARK signal following DMSO or 10 µM CCCP treatment (*p = 0.0248; unpaired t-test, two-tailed).

For all images, scale bars represent 10 µm. Bar graphs display mean ± standard deviation.

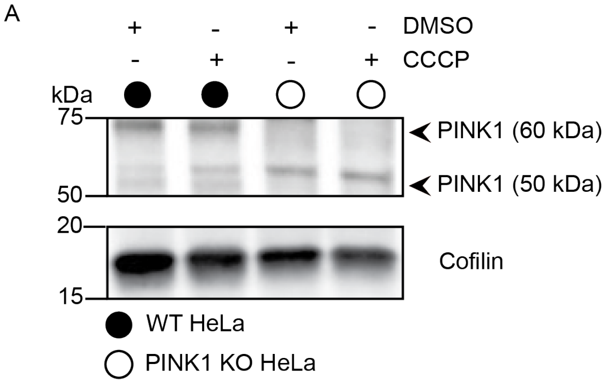


**Supplementary Figure 2. Confirmation of PINK1 knock out in HeLa cells.**

A. Western blot of WT and PINK1 KO HeLa cells treated with DMSO or 10 μM CCCP.
